# The timid invasion

**DOI:** 10.1101/2022.06.16.494985

**Authors:** Jana A. Eccard, Valeria Mazza, Celia Holland, Peter Stuart

## Abstract

In biological invasion processes animal behavior moderates the success of invasive species, and the native fauna’s ability to adapt. The importance of behavioral syndromes and flexibility of invading species for invasion success remains debated. We investigated behavior of rodents, the bank vole (*Myodes glareolus*) currently invading Ireland, and the native wood mouse (*Apodemus sylvaticus*) that declines with vole invasion, at replicated sites at pre-invasion, edge, and source of the invasion. We found that individual rodents varied consistently in risk-taking behaviors, and mice had not adapted to the presence of invasive voles. Voles at the invasion edge were more careful but also more flexible compared to voles the invasion source. The ability to develop timid and flexible phenotypes may contribute to the invasion success of rodents worldwide.

## Main Text

Behavior is a flexible trait that allows animals to cope with novel environments and challenges (1), allowing, for example, adjustments to habitats in which a species had not evolved (2, 3). Recent growth in the study of inter-individual variation in behavior (4, 5, 6), however, revealed that behavior is not completely flexible (5) and that individuals differ consistently within a population (7). Further, this consistent variation has fitness consequences (8, 9). For the success of biological invasions of animals, two contrasting concepts have been suggested: behavioral invasion syndromes and behavioral flexibility (10). Consistent inter-individual differences in behavior, with sets of traits that occur together, are called behavioral syndromes (5), and traits increasing invasion success include high dispersal tendency, high foraging efficiency, and risk-taking in new environments (for review 11). On the other hand, behavioral flexibility, i.e. the ability to respond quickly to altered environmental conditions, is believed to have facilitated successful invasions. Meanwhile, flexibility is often inferred indirectly by comparisons of relative brain size (12, 13) or by habitat plasticity (14). Here we aim to disentangle the relative contribution of consistent inter-individual variation in behavior (repeatability) and behavioral flexibility to biological invasions, by investigating a currently expanding population of rodents with repeated, behavioral tests which allow the quantification of both mechanisms.

Invasions of animals to previously unoccupied areas can be distinguished into two separate spatiotemporal processes: expansion and settlement, during which selective pressures on traits related to dispersal e.g. morphological trait distribution within the population, can shift (15, 16). To understand how invasions occur and which candidate traits may favor expansion in a non-native environment, we need to monitor an expansion in real time. An ongoing rodent invasion of Ireland by a continental vole species (17), that started a century ago and currently covering more than half of Ireland, offers a unique opportunity. We investigated rodent behavior of both the invasive bank vole (*Myodes glareolus*) and the single native small rodent species, the wood mouse (*Apodemus sylvaticus*) at replicated populations in different invasion zones, testing 450 rodents in 535 behavioral tests. At the invasion source voles had been present for 80-100 years (17) and co-exist with low numbers of mice; at the invasion edge voles appeared mice and voles coexist both in high numbers; and pre-invasion mice were found in high numbers (Fig. 1A, and (18) data and methods available as supplementary materials (tables S1-S5) at the Science website).

**Figure 1.**
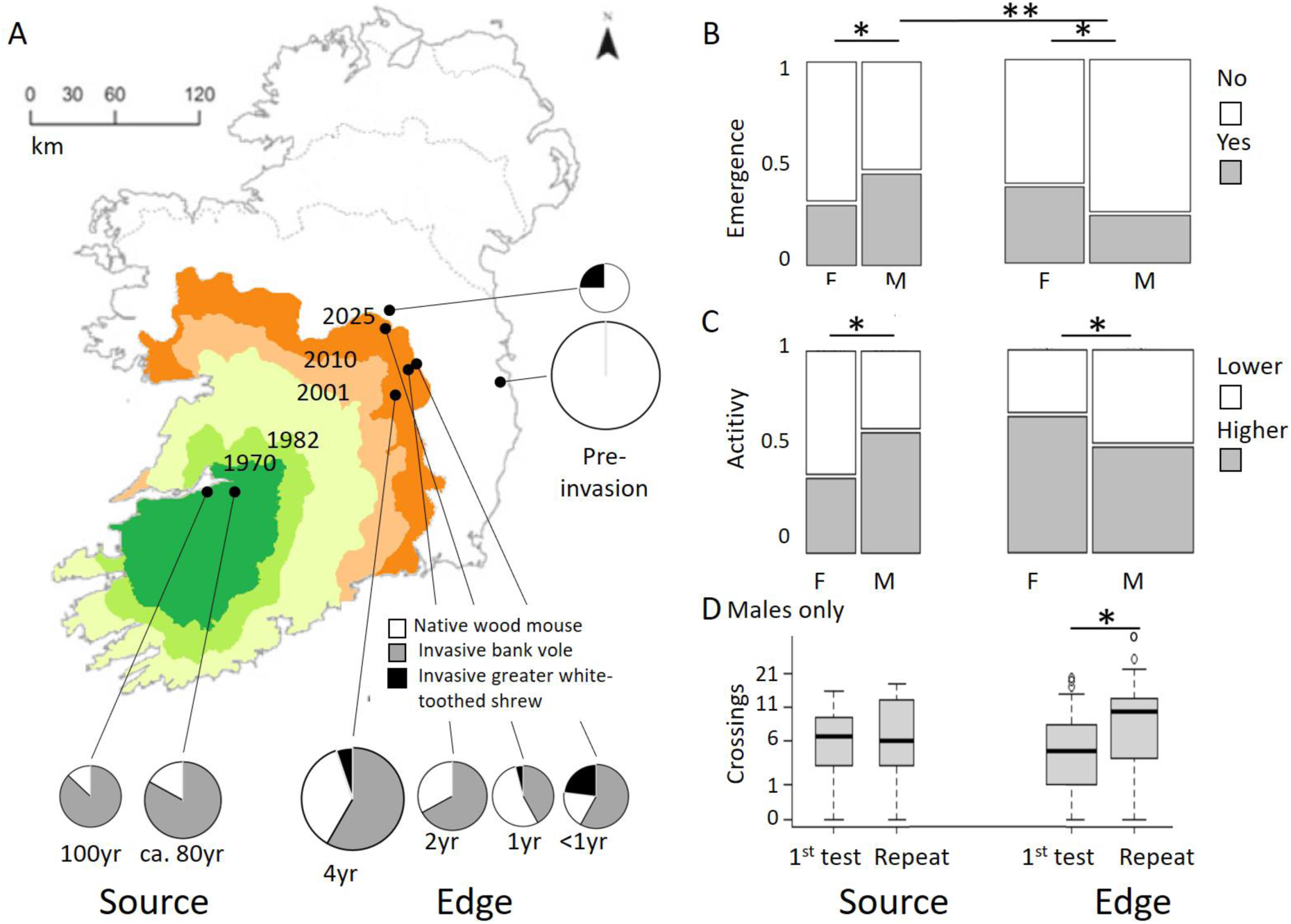
A. Invasion history of bank voles introduced a century ago to Ireland (dark green area). Boundaries between colored invasion ranges according to White et al. 2010. Sampling populations: black dots, size of corresponding pie chart indicates population density during the study, ranging from 9 (smallest) to 53 (largest), small rodents per 100 trap nights. Shading of pie chart depicts local species composition at capture. B-D Risk taking behavior and activity of invasive bank voles in source and edge zone of the invasion. (B) Emergence within 5 minutes in a dark-light test; (C) activity type in an open field arena, % activity peaked bimodally at <10% or >90% activity and was therefore analyzed as a bimodal variable (lower activity, higher activity with cut-off at 50%); and (D) risk-taking (self-exposure) in an open field arena indicated by crossing of the center. Shown are interactive effects of invasion zone*sex of the tested animal (A, B) and behavioral flexibility of males, i.e. first versus repeated testing (C, statistic Tables S2-4). Width of bars and boxes indicate relative sample size.

We focused on two aspects of behavior which are likely to influence individuals’ success during the various stages of colonization (*sensu* 19), i.e. the amount of exploration activity in an unfamiliar environment, and the propensity to take risks, or boldness. Bolder individuals are expected to reap greater rewards in terms of mates and/or resources, and thus to have better chances to navigate novel or potentially hostile environments (e.g. 2; 19). Inter-individual variation in exploration activity predicts dispersal tendency and space use in several species (e.g. 20), and is also suggested to facilitate range expansion in non-native habitats (e.g. 21). We hypothesized that individuals at the edge of the expansion would differ in the mean expression of personality traits related to risk-taking and exploration, as well as in the level of flexibility of those traits. We predicted that (a) voles at the expanding edge of the invasion would take more risks compared to individuals at the source of the invasion, and (b) that voles at the expansion edge would be behaviorally more flexible and habituate faster to the novel test situation compared to those from the source, as they are more likely to experience unpredictable conditions and unsuitable habitats. Since biological invasions have the deepest impact in terms of effects on native fauna/biodiversity, we additionally investigated whether the native wood mice would show behavioral adjustments in response to the presence of bank voles. We investigated the behavior of mice in the same source (two site replicates) and edge populations (4 site replicates) where bank voles were investigated, as well as in two pre-invasion sites, which were still vole-free. We predicted, since mice numbers decline with persistence of voles (22), that mice behavior did not mitigate effects of competition with voles and (c) mice behavior does not differ among invasion zones.

Animals were captured overnight, tested for risk-taking and exploration, then sexed and weighed, marked by an individual fur cut, and photographed for later identification. Sites were visited for two trapping sessions within 4 days, and re-visited after 2-4 weeks to allow repeated testing on recaptured individuals. Population structure of both species (distribution of age classes, sexes, survival rates (i.e. recapture), body weight) did not vary among zones of the invasion (mixed effect models based on individual captures, including the individual ID and the research visit ID, Table S2). We quantified risk taking propensity and activity of 225 voles and 189 adult mice in 535 behavioral tests, using personality tests established for behavioral phenotyping of small rodents under field conditions (20). Not all data could be used in all analyses, since some mice managed to escape before marking or sexing (Data S1). We assessed boldness by measuring individuals’ willingness to emerge from a dark covered shelter into an open, illuminated, and empty arena, which is perceived as dangerous for a rodent species with a plethora of predators, within a 5min observation window. We quantified exploration activity by measuring the proportion of time each individual spent moving in the novel space of the arena, and also how often they crossed into its central part, which should be perceived as being even more exposed and potentially dangerous. We found that for both rodent species the recorded behavior varied consistently among individuals, in relation to the variance measured among conspecifics (see repeatability scores for emergence, exposure and activity in table S1), allowing us to investigate adaptations at the phenotypic, individual level.

At the edge of the invasion, male bank voles were shyer than conspecific males at the source of the invasion, as indicated by a lower proportion (24%) emerging from the dark covered shelter into the open test arena, compared to individuals from the source of the invasion (49%, mixed model with binomial distribution, the invasion zone explained the behavioral variables in an interaction with sex of the tested animal with P < 0.002, Tables S3a and Post-hoc tests Table S4). Further, activity of males during the test was lower than of females at the edge of the invasion, while males were more active than females at the source of invasion. While male fish and birds appear to be more aggressive at the invasion edge (23, 24) the invasive rodents observed here were less bold at the edge, compared to conspecifics living in the source of the invasion. Rodents are a highly predated-upon taxon, and have evolved to a rather cryptic lifestyle avoiding open spaces and dangerous daytimes, which seem to favor increased cryptic and timid behavior in an invasion processes. Many small rodent populations undergo population fluctuations with dramatic regional population crashes at times (e.g. 25), and may thus have a pre-adaptation to re-colonization processes.

At the expansion edge, male voles were more flexible than at the source, as indicated by stronger habituation to repeated testing. With repeats, male voles at the edge increased their exploration of the risky area (number of crossings: z = 3.5, p > 0.001), while males in the source population did not (all z < 0.8, p > 0.407, interaction of zone and habituation Table S3a). Activity also increased with repeated testing at the invasion edge, (Figure 1D, Table S2 and Post-hoc tests Table S4, z = 2.1, p = 0.038), but not at the source (z = 1.3, p = 0.185, interaction of factors Table S3a). Rodents are able to adjust to man-made changes in their environment (e.g. urbanization, 3, 26) and challenging environments, by increasing their behavioral plasticity both at the genotypic and phenotypic level. Genetic comparisons between established range and edge population of bank voles already suggested adaptations potentially coding for behavior and sexual differentiation (17), which was confirmed at the phenotypic level by the present study. Furthermore, behavioral plasticity is related to risk taking behavior in bank voles (27) and our results show that both are relevant traits in dispersal processes. The ability of a species to adjust to novel challenges is likely to contribute to its ultimate success in novel environments.

Sex differences operated in the opposite direction in the two zones: in the expansion zones, males were more risk-averse and less active compared to females, while at the source, males were more risk-prone and more active (GLMM, interaction of zone*sex: emergence: chi2=7.2, p= 0.007; activity chi^2^ = 8.7, p = 0.003, nr. of center crossings chi^2^=11.4, p < 0.001, Fig. 1B, Table S3a, Post-hoc tests Table S4). Behavior of bank vole females did not differ between zones (z = 1.0, p = 0.320, Post-hoc tables ESM2b). In rodents, males are the dispersing sex. Avoiding risks may be a trait that is under strong selection in the expansion process of a population while it disperses into areas void of conspecifics, and may thus be best measurable in the dispersing sex.

Wood mice behavior did not differ between the invasion zones of bank voles or the pre-invasion zone (Table S3b). At every zone, the individual mice’s test experience (repeated testing) increased the probability to emerge, number of crossings and activity (Table S3b). Our results on mice strengthen the results of parallel tests on voles, showing that there are no effects of environmental or temporal variables except geographic location (i.e. invasion zone) in voles, which would have affected both species. Population sizes of wood mice are negatively related to the increasing populations of invading bank voles (22). A lack of behavioral differentiation by mice in response to voles’ presence allows several interpretations. Wood mice and bank voles are sympatric throughout Europe, except in Ireland. Mice arrived first in Ireland a millennium ago (28), and if competitive release was not accompanied by behavioral changes it could not be reversed by the re-appearance of the competing species. Furthermore, risk-taking behavior, as measured here, may rather be affected by predation risk than by competition. In addition, crepuscular and nocturnal mice may not reveal behavioral adaptations when studied during bright daylight. Lastly, voles and mice are partially segregated in their daily activity patterns, potentially decreasing strong behavioral interference. Numerical competitive effects on nocturnal mice may mainly be due to indirect resource depletion by voles, which consume similar food resources but are able to deplete resources at both day and night.

To conclude, our study confirmed the importance of both a behavioral invasion syndrome at the edge of the vole’s invasion range, but also the increase of behavioral flexibility of individuals at the expansion edge. Invasive bank voles in Ireland were more cautious at the invasion edges of the species distribution and habituated faster in repeated tests compared to voles at the source of expansion. This rather timid dispersal strategy of a rodent at the expansion edge may explain why rodents worldwide are a highly successful, invasive taxon. Results may have implications for the management of invasive species, requiring different measures at expanding populations compared to established ranges.

## Acknowledgements

We thank our field helpers Paul Mosnier, Ciaran O’Ciuv and Matthew Quinn, the site managers, and TCD Zoology department for support.

## APPENDIX

### Materials and Methods

#### Study locations in source and edge sites

Bank vole populations at the expansion edge were studied in summer 2019 when the expansion edge had migrated 200-250 km east of the source population, at 2 sites near the source, 4 near the 2019 distribution edge, and 2 pre-invasion sites (Article Figure 1A). Sampling locations in forest fragments (size 3ha-100ha) were selected based on Stuart et al. (29). Rodents were trapped in single-capture Longworth traps (Penlon Ltd., Oxford, UK) which were pre-baited with seeds for 2 nights prior to trapping.

#### Trapping procedures and recapture rates

At each site, 24 traps were set in lines with 10 m spacing between them. Traps were set over night and checked in the morning. Trapping and testing were not conducted during rainy days. Animals were kept in their traps in the shade until testing, with additional food and water sources.

We captured invasive bank voles, native wood mice, and a new invasive species (22), the greater white toothed shrew (*Crocidura russula*, proportions of captured species per site are shown in Figure 1A). With the focus of the study on behavior, we report the population background of the sampled animals here as part of the methods. As in Stuart et al. (29), the proportion of voles relative to mice were higher at the source sites than at the expansion edge (Fig. 1A). Across all sites, we captured and tested 196 individual bank voles, of which 67% were adults and 54% of adults were male. Recapture rate of voles was 35% of animals during the same visit, and 27% during the next visit to the same site. With mice, we captured 237 individuals across sites, however a third of the agile mice escaped unhandled and unmarked during or after the test procedure. Excluding these captures (and also excluding recapture status of unmarked animals at a specific site after the first unmarked animal had escaped), 80% of identified mice were adult and 58% of adults were male. 32% of animals were identified recaptures during the first and 42% during the second visit. Rodent captures per trap (mice and voles) and demography of populations (sex, age, body weight, probability of recapture, i.e. a surrogate of population turnover rates) of each species were independent of the expansion zone (Tables S2, generalized linear mixed models based on captures, using animal ID and trapping session as random variables, z < 0.5, P > 0.61 in voles (2 zones); z < 1.1, P > 0.251 in mice (3 zones)).

#### Assessing risk-taking and activity

For behavioral testing, we adopted two standardized behavioral tests that are commonly used in animal personality studies of small mammals to assess boldness and spatial exploration in a novel environment. The set-up combined the dark-light test (30) and the open-field test (31), and was adjusted to be executed under field conditions without prior handling of animals (e.g. 20, 3). Test arenas were always set in shady locations or under canvas roofs to avoid direct sunlight, shade patterns and open sky above the arena. At the end of each test, the arenas were cleaned with ethanol 70%. All tests were conducted between 1000 and 1800, under natural light conditions.

##### Dark-light test

An opaque plastic tunnel (15 cm diameter, 30 cm long) was connected to an opening in the circular arena at ground level, blocked by a wooden door. Directly before the tests, the animal was transferred from the trap into a shuttle tube (15 cm diameter, 30 cm long, open at one side and blocked by a cylindrical sponge of the same diameter on the other). The shuttle tube with the animal inside was then inserted into the plastic tunnel. After 1 minute of acclimatization to the dark tunnel, the door leading into the arena was opened and we measured the rodent’s latency to emerge (emergence) from the dark tunnel into the arena with its full body (without the tail). If animals did not emerge from the tunnel within 5 min (66% of tests in voles and 24% in mice), we slowly reduced the space in the tube by pushing in the soft, cylindrical sponge in order to gently displace the animal without touching it.

##### Open field test

We used round, plastic, foldable arenas (1.20 m diameter, 40 cm high) as portable test set-ups. Arenas were covered with mesh to prevent mice jumping out. The open field test started as soon as the animals entered the arena from the dark light test with its full body. The door leading to the dark tube was closed quietly by the observer, or, if the animal was pushed out, the tube remained blocked by the sponge. The floor of the arena was divided into a 15-cm wide zone along the wall (known to be perceived as safe by the rodent), and the open center of the arena (perceived as dangerous). For 5 minutes after emergence, we counted the number of 10 sec intervals spent moving in the arena (activity), the number of crossings into the central part of the arena with the whole body (crossings). After testing, animals were removed from the arena by offering a hiding place, either by pulling back the sponge in the tube or by placing their familiar trap in front of them.

#### Data cleaning and statistical analyses

##### Modelling individual behavior in the different expansion zones

We investigated inter-individual differences in boldness and activity between rodents at the expansion edge and at the established source area using generalized linear mixed models (GLMM with R package lme4). We added expansion zone (2 zones for bank voles, 3 zones for mice (Fig. 1A)), sex (male vs female), and test repeat (first test vs repeated tests to account for habituation to testing conditions) as fixed effects to model behavioral variables. We used a combination of siteID and visitID (23 combinations) as random effect, to account for a) temporal (daytime, weather, season, involved team members) and spatial (sites) variation among testing occasions, and b) dependence of tested individuals from the same place and time. We used animal ID as a second random factor to control for repeated testing of the same individuals (a third of the tests were repeats). Initial models included two-way interactions of the main effects, which were removed if not significant. We report log-likelihood tests (Wald’s chi^2^) in the text and show effect sizes in tables and provide R^2^ to indicate variation explained without (marginal) and with (conditional) random factors (32).

Since juvenile mice did not move in either round, we stopped testing them and did not analyze the data initially collected on juveniles. We used all available mice data (including animals that escaped before ageing, sexing and individual identification) for a model comparing between zones. We conducted additional analysis including zone and individual information (sex, repeated testing) with a reduced data set on identified mice only (see table S3 for sample sizes for different variables).

In the vole dataset, variables were zero inflated (29% of voles never crossed into the center of the arena) or bimodal (voles left the dark space either immediately or not at all during the 5 minutes; activity in the open field had peaks at <10% or >90% activity) and were therefore analyzed as binomials. Within the trials where animals had actually crossed into the center of the arena, we also analyzed the number of crossings.

##### Assessing the repeatability of inter-individual variation in behavior

As a precondition to our comparisons of individual behavior among invasion zones, we first challenged the assumption that behavioral traits are consistent within individuals and are thus a useful representation of an animal’s behavioral type or personality (6). This precondition was confirmed as indicated by the repeatability of measured trait values over time (33), (Table S1). Repeatability is a population-specific metric used to quantify among-individual phenotypic differences across time or contexts relative to the variation in the population. We calculated R with the R package *rpt*R (34), using individual ID as a random factor in generalized linear mixed models (G)LMM including the first vs repeated testing (33). The boldness measures, i.e. the emergence in the dark-light test, and the number of crossings of the open field arena were repeatable (in tendency for emergence in mice), while activity was not. For repeatability analysis of mice we only used data of animals with confirmed recapture status (first capture or recaptured).

**Table S1:**
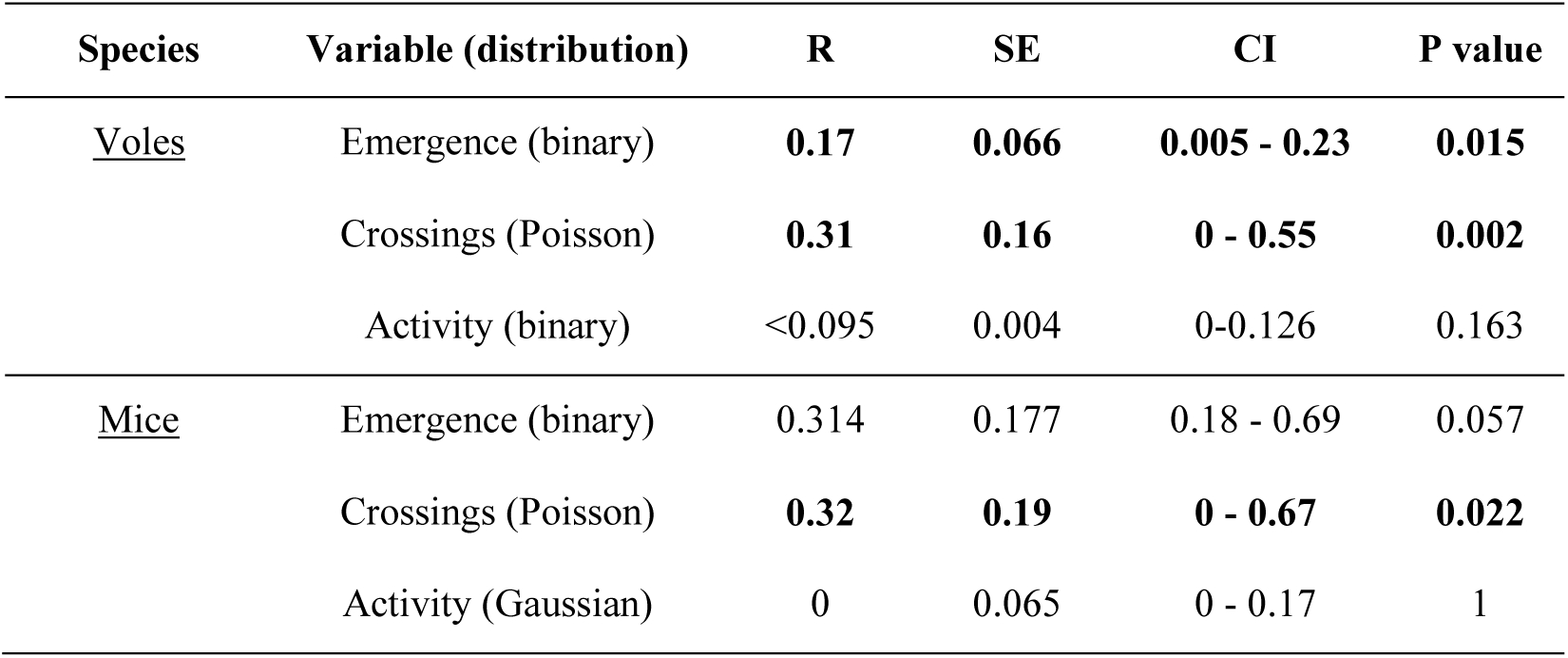
Repeatability. (R) estimates, standard errors (SE) and 95% confidence intervals across test repeats for 189 individual invasive bank voles (*Myodes glareolus*, 314 tests) and 82 individual native wood mice (*Apodemus sylvaticus*, 111 tests). The distribution of each behavioral variable, as specified in the models, is reported in (). Significant effects are marked in bold font.

#### Ethical note

Ethical approval for this work was obtained from Trinity College Dublin, Ireland, Animal Research and Ethics Committee (Ref. 130219). Site use was permitted by the respective county or local park authorities. Behavioral experiments were performed in accordance with all applicable international, national and/or institutional guidelines for the use of animals, including the ASAB/ABS Guidelines for the use of animals in research. We took great care in ensuring the animals’ welfare throughout the experimental procedure and afterwards. Traps contained hay as bedding, bird food as bait, and a piece of vegetable as water supply. Traps were checked after 8 hours, and refilled with food and vegetable if occupied. Occupied traps were kept in the shade until testing. Handling was done with care and only on first capture, individual markings were based on non-invasive fur cutting. After testing, animals were released at the point of capture.

### Supplementary Text

#### Extended technical descriptions of results: Sex-specific behavior at the expansion edge

At the expanding edge of the invasion, bank vole males were more risk-averse than females, while in the source population there was no difference (29%; GLMM, interaction zone*sex log-likelihood test: chi = 7.2, p = 0.008, table simple effects, Figure 1B). Bank vole males at the expansion edge were less active then females, while males’ activity was higher than females at the source (interaction zone*sex, chi = 8.7, p = 0.003, Figure 1C). At the established range, sex differences were expressed in the opposite direction: a larger proportion of males were bolder and more active (post-hocs, z > 2.0, p = 0.044), while at the expansion edge, males were more risk-averse and less active (z <-2.2, p < 0.029) compared to the females in the respective zone (interaction of zone*sex: emergence: chi2=7.2, p= 0.007; activity chi2 = 8.7, p = 0.003, number of crossings into the center of the arena chi=11.4, p < 0.001, Fig. 1, Table ESM2a).

At the expanding edge of the invasion, males were least likely to emerge from the dark into the open test arena (24% emerged), compared to males at the established source of the invasion (49%), and compared to the females at the edge (38%), while in the source population, male emergence was similar to females (29%; GLMM, interaction zone*sex log-likelihood test: chi = 7.2, p = 0.008, table simple effects, Figure 1B). Furthermore, edge males’ activity in the open field test was lower compared to that of females at the edge, while again males’ activity was higher than females at the source (interaction zone*sex, chi = 8.7, p = 0.003, Figure 1C). Among the animals which crossed into the center of the open field, exposed themselves (crossing the open center), males at the edge increased the number of crossings with repeated testing, while males from source populations and females from either zone did not change their behavior (interactions zone*sex chi = 11.4 p < 0.001, zone*repeat chi = 5.6, p = 0.017, Table xx, males: Figure 1D, females: median 5 crossings, range 0-20 crossings).

Once in the open field, the probability of an animal to cross into the center of the arena was higher when an individual was tested repeatedly (chi = 8.2, p = 0.003, 59% in first test, 74% in repeated tests) and higher for females at the edge (74%) than at the source (43%), while for males, zones did not differ (67%, 68%, interaction zone*sex chi = 6.8, p = 0.009).

At the established range, a larger proportion of males emerged and males were more active (post-hocs, z > 2.0, p = 0.044), while at the edges, a smaller proportion of males emerged and less were active (z <-2.2, p < 0.029) compared to the females in the respective zone (interaction of zone*sex: emergence: chi2=7.2, p= 0.007; activity chi2 = 8.7, p = 0.003, nr. of center crossings chi=11.4, p < 0.001, Fig. 1, Table ESM2a). Activity of males did not differ among zones (simple effects, emergence z = −0.7, p = 0.499) and behavior of females did not differ between zones (z = 1.0, p = 0.320, Posthoc tables ESM2b).

Habituation rates of bank voles were higher at the edge compared to established ranges. At the edge of the distribution animals were more likely to emerge in the repeated test compared to their first test (post hoc table, z = 2.1, p = 0.038), while this was not the case at the source (z = 1.3, p = 0.185, interaction of factors table ESM2a). At the edge, males increased the number of crossings into the center of the arena from the first to repeated testing (z=3.5, p > 0.001), while this was neither the case for females in any of the zones, nor for males in the established range (all three z < 0.8, p > 0.407)

**Table S2a.**
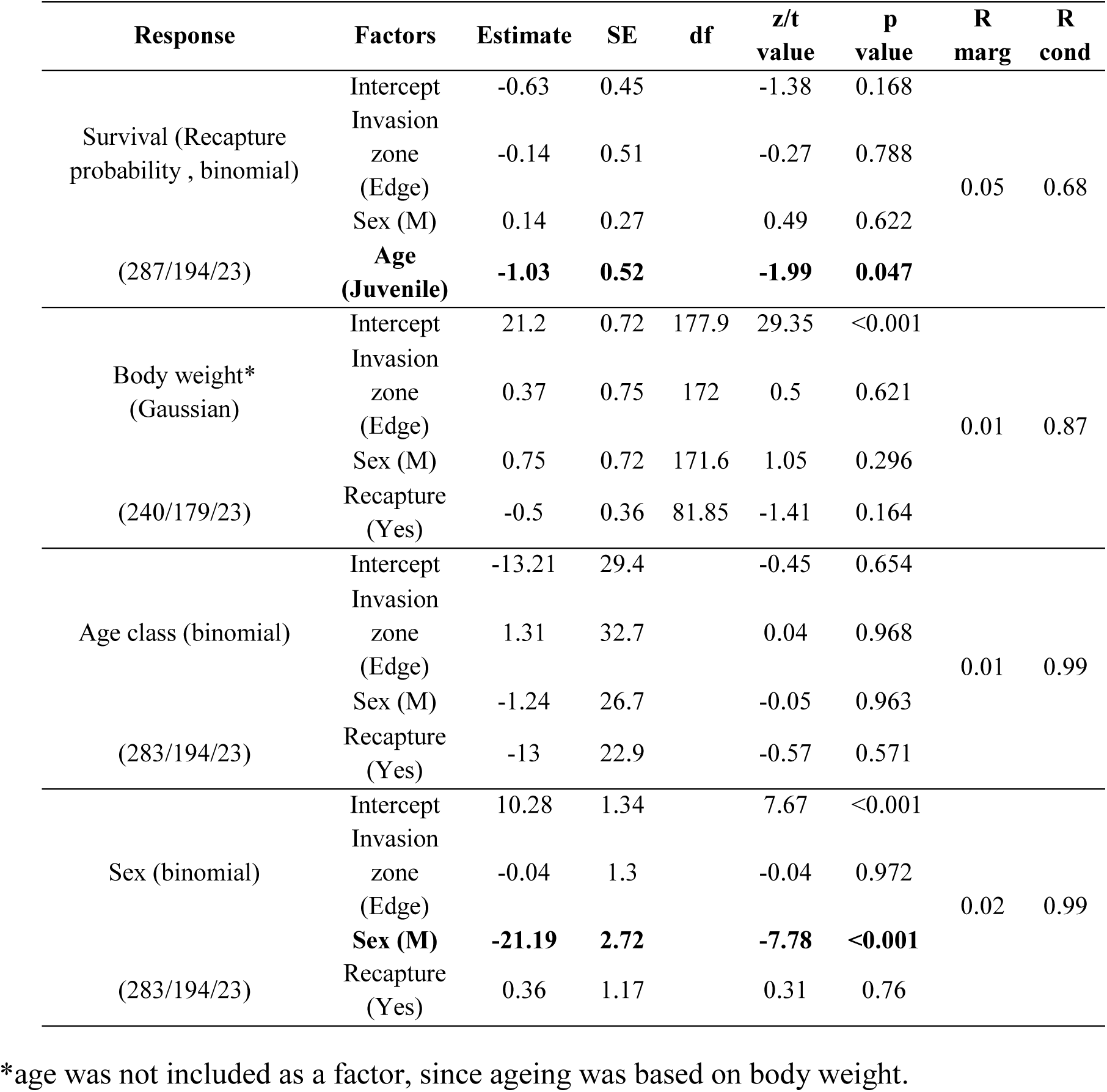
Population structure bank voles. Survival probability (recapture), body weight, age class and sex distribution of captures of invasive bank voles in Ireland, analyzed with generalized linear mixed models (GLMM. package lme4). Brackets indicate nested sample sizes (nr. of tests/nr. of individuals/number of trapping ID (site*date)). Trapping ID was included to account for potential non-independence of captures at the same site on the same day. Population structure did not differ among expansion zones. Effect sizes and directions refer to expansion edge relative to source populations (zone), males relative to females (sex), juveniles to adults (age), recapture relative first capture (capt). Statistical models included individual ID and test occasions ID as random effect. R_marg_ and R_cond_ indicate variation explained without (marginal) and with (conditional) random factors. Across sites, recapture probability (a) was higher for adults than for juveniles, and (d) we had captured more adult males than females and more juvenile females than males. Significant effects are marked in bold font.

**Table S2b.** Population structure wood mice. Population structure and survival probability of native wood mice populations in Ireland analyzed with generalized linear mixed models (GLMM. package lme4) based on captures. Brackets indicate sample sizes (nr. of captures/of individuals/of test occasions). Populations did not differ among expansion zones. Effect sizes and direction refer to expansion edge and pre-invasion zone relative to source populations (zone), males to females (sex), juveniles to adults (age), recapture relative first capture (capt). Statistical models included individual ID and test occasions ID as random effect, Rmarg and Rcond indicate variation explained without (marginal) and with (conditional) random effects. A model for age class did not converge.

| Response | Factors | Estimate | SE | df | z value/t value | p value | R<sub>mar</sub> | R<sub>cond</sub> |
| --- | --- | --- | --- | --- | --- | --- | --- | --- |
| Survival<br>(Recapture probability binomial)<br>(155/102/28) | Intercept | -1.6 | 1.6 |  | -1.02 | 0.308 |  |  |
|  | Invasion zone (Edge) | -2.04 | 1.78 |  | 1.14 | 0.253 | 0.07 | 0.87 |
|  | Invasion zone (Pre-invasion) | -0.16 | 2 |  | -0.08 | 0.935 |  |  |
| Body weight<br>(Gaussian)<br>(103/79/26) | Intercept | 22.52 | 2.19 | 80.5 | 10.2 | <0.001 |  |  |
|  | Invasion zone (Edge) | 1.1 | 2.31 | 75.97 | 0.5 | 0.431 |  |  |
|  | Invasion zone (Pre-invasion) | -1.1 | 2.4 | 66.75 | 0.05 | 0.65 | 0.33 | 0.84 |
|  | Age<sup>§</sup> (Juvenile) | **-8.5** | **1.27** | **94.73** | **-6.7** | **<0.001** |  |  |
|  | Sex (M) | -0.98 | 0.99 | 89.15 | -1.03 | 0.306 |  |  |
|  | Recapture (Yes) | -0.15 | 0.75 | 35.3 | 0.19 | 0.845 |  |  |
| Sex<br>(binomial)<br>(124/89/28) | Intercept | 9.6 | 3.03 |  | 3.18 | 0.001 |  |  |
|  | Invasion zone (Edge) | 0.92 | 3.12 |  | 0.29 | 0.769 |  |  |
|  | Invasion zone (Pre-invasion) | -0.63 | 3.06 |  | -0.21 | 0.838 | 0.01 | 0.99 |
|  | Age (Juvenile) | 1.65 | 2.27 |  | 0.72 | 0.466 |  |  |
|  | Recapture (Yes) | -0.39 | 1.37 |  | -0.29 | 0.774 |  |  |

<sup>§</sup>Age was included since ageing was based on fur structure

**Table S3a.** Bank voles: Full models of behavioral variables. emergence, crossings and activity compared between two main invasion zones (source and edge), sexes (M and F), and habituation to the test (recapture Yes/No); analyzed with a generalized linear mixed models (GLMM. package lme4). Brackets indicate sample sizes (nr. of tests/rando effect individuals/random effect test occasions). Given are effect sizes (Estimate) and standard error (SE) and sample sizes; R_marg_ and R_cond_ indicate variation explained without (marginal) and with (conditional) random effects. Significant effects are marked in bold font.

|  | Factors | Estimate | SE | df | z/ t value | p value | R marg | R cond |
| --- | --- | --- | --- | --- | --- | --- | --- | --- |
| Emergence<br>(binomial)<br>(291/202/26) | Intercept | -1.54 | 0.53 |  | -2.92 | 0.003 |  |  |
|  | Invasion zone (Edge) | 0.61 | 0.59 |  | 1.04 | 0.298 |  |  |
|  | **Sex (M)** | **1.37** | **0.58** |  | **2.35** | **0.019** | 0.17 | 0.59 |
|  | Recapture (Yes) | 0.66 | 0.35 |  | 1.88 | 0.06 |  |  |
|  | **Invasion zone (Edge)\*Sex (M)** | **-2.33** | **0.77** |  | **-3.04** | **0.002** |  |  |
| Crossings<br>(Poisson)<br>(203/150/24) | Intercept | 1.53 | 0.23 |  | 7.15 | <0.001 |  |  |
|  | Invasion zone (Edge) | 0.3 | 2423 |  | 1.25 | 0.221 |  |  |
|  | Recapture (Yes) | -0.19 | 0.16 |  | -1.21 | 0.226 |  |  |
|  | **Sex (M)** | **0.59** | **0.18** |  | **3.25** | **0.001** | 0.02 | 0.33 |
|  | **Invasion zone (Edge)\*Sex (M)** | **-0.79** | **0.23** |  | **-3.28** | **<0.001** |  |  |
|  | **Recapture (Yes)\*Sex (M)** | **0.45** | **0.19** |  | **2.38** | **0.017** |  |  |
| Activity<br>(binomial)<br>(290/199/26) | Intercept | -0.6 | 0.44 |  | -1.33 | 0.18 |  |  |
|  | **Invasion zone (Edge)** | **1.29** | **0.53** |  | **2.43** | **0.015** |  |  |
|  | **Sex (M)** | **0.91** | **0.43** |  | **2.11** | **0.035** | 0.31 | 0.53 |
|  | Recapture (Yes) | 0.48 | 0.28 |  | 1.74 | 0.082 |  |  |
|  | **Invasion zone (Edge)\*Sex (M)** | **-1.64** | **0.55** |  | **-2.99** | **0.003** |  |  |

**Table S3b.**
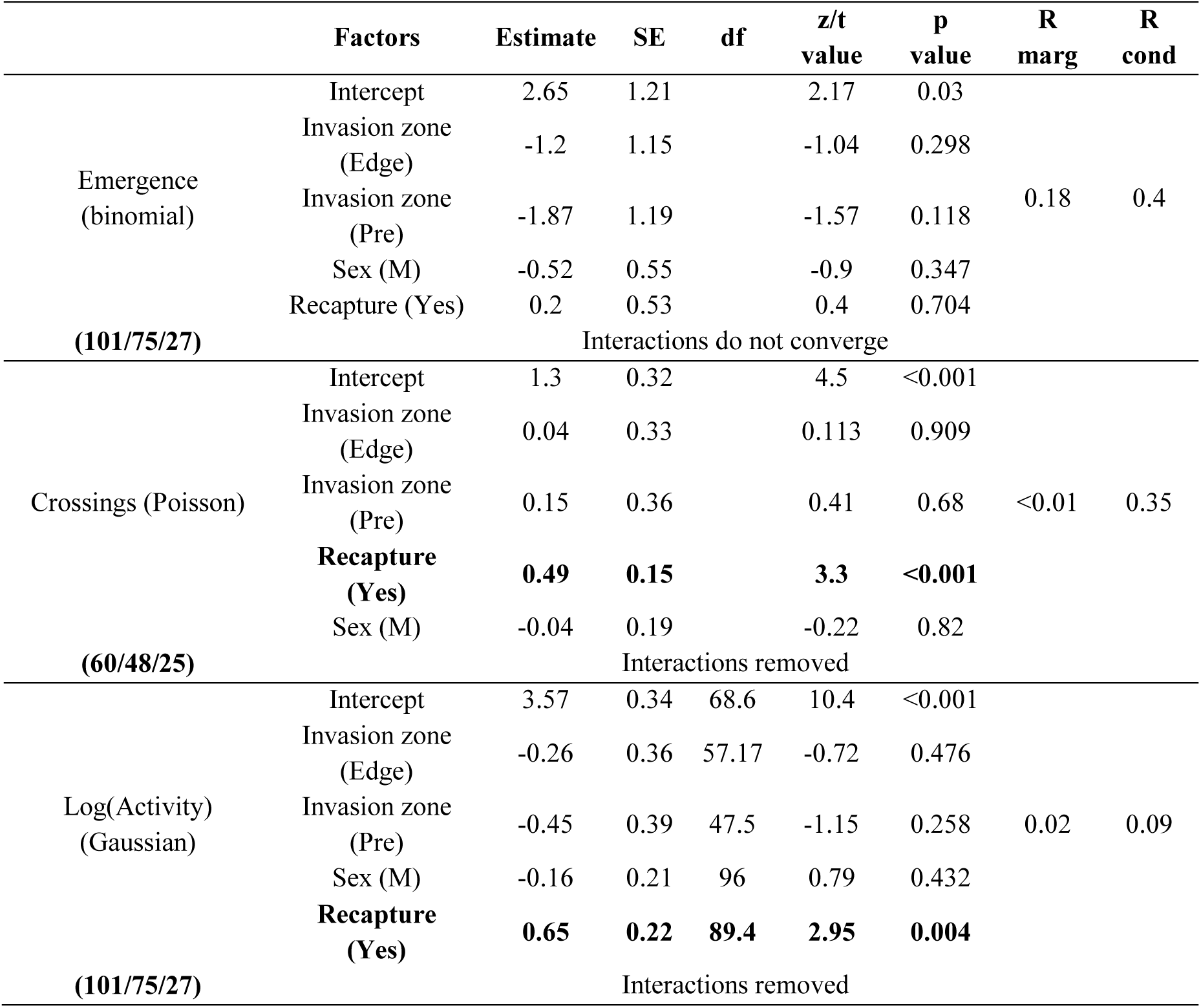
Wood mice: Full models of behavioral variables. emergence, crossings and compared among three bank vole invasion zones (source, edge, pre-invasion), sexes (M and F), and habituation to the test (recapture Yes/No); analyzed with a generalized linear mixed models (GLMM. package lme4). Brackets indicate sample sizes (nr. of tests/random effect individuals/random effect test occasions). Given are effect sizes (Estimate) and standard error (SE) and sample sizes; R_marg_ and R_cond_ indicate variation explained without (marginal) and with (conditional) random effects. Significant effects are marked in bold. We included factors analogue to the analysis of bank vole behavior. Reduction of models sometimes allowed for larger sample sizes to analyze the remaining factors, however is not shown since estimates remained largely unchanged. Significant effects are marked in bold font.

**Table S4.** Bank vole behavior - Disentangling interactions from Table S3a: Simple-Effects within respective subsets analyzed with generalized linear mixed models (GLMM) as in Table S3. Brackets indicate sample sizes nested within random effects (nr. of tests/nr. of individuals /nr. of test occasions). Given are the standard error (SE). Marginal and conditional R2 values were similar as given for full models (Table 2a). Interactions were removed if not significant in any of the post-hoc analyses. Significant effects are marked in bold font.

|  | Data subset | Factors | Estimate | SE | z/t value | p value |
| --- | --- | --- | --- | --- | --- | --- |
| Emergence (binomial) | **Males**<br>(162/103/21) | Intercept | -0.3 | 0.43 | -0.71 | 0.478 |
|  |  | Invasion zone (Edge) | **-1.56** | **0.61** | **-2.58** | **0.010** |
|  |  | Recapture (Yes) | 0.7 | 0.45 | 1.56 | 0.118 |
|  | **Females**<br>(129/99/21) | Intercept | -1.44 | 0.67 | -2.14 | 0.032 |
|  |  | Invasion zone (Edge) | 0.41 | 0.61 | 0.67 | 0.501 |
|  |  | Recapture (Yes) | 0.69 | 0.6 | 1.14 | 0.253 |
|  | **Edge**<br>**179/125/18)** | Intercept | -0.96 | 0.39 | -2.44 | 0.015 |
|  |  | Sex (M) | **-1.01** | **0.46** | **-2.21** | **0.027** |
|  |  | Recapture (Yes) | **0.87** | **0.43** | **2.02** | **0.044** |
|  | **Source**<br>(112/79/15) | Intercept | -1.84 | 0.81 | -2.29 | 0.022 |
|  |  | Sex (M) | **1.78** | **0.7** | **2.55** | **0.011** |
|  |  | Recapture (Yes) | 0.24 | 0.54 | 0.44 | 0.658 |
| Crossings (poisson) | **Males**<br>(109/75/20) | Intercept | 1.45 | 0.24 | 5.96 | <0.001 |
|  |  | Invasion zone (Edge) | -0.04 | 0.3 | -0.14 | 0.891 |
|  |  | Recapture (Yes) | 0.24 | 0.22 | 1.06 | 0.289 |
|  |  | Invasion zone (Edge)*Recapture (Yes) | 0.31 | 0.27 | 1.13 | 0.257 |
|  | **Females**<br>(80/66/20) | Intercept | 1.5 | 0.2 | 7.47 | <0.001 |
|  |  | Invasion zone (Edge) | 0.29 | 0.23 | 1.3 | 0.193 |
|  |  | Recapture (Yes) | -0.08 | 0.27 | -0.28 | 0.78 |
|  |  | Invasion zone (Edge)*Recapture (Yes) | 0.18 | 0.3 | 0.59 | 0.557 |
|  | **Edge**<br>(126/89/18) | Intercept | 1.82 | 0.14 | 12.84 | <0.001 |
|  |  | Sex (M) | -0.15 | 0.16 | -0.98 | 0.329 |
|  |  | Recapture (Yes) | 0.07 | 0.14 | 0.48 | 0.629 |
|  |  | **Sex (M):Recapture (Yes)** | **0.39** | **0.19** | **2.1** | **0.036** |
|  | **Source**<br>(63/53/6) | Intercept | 1.51 | 0.18 | 8.42 | <0.001 |
|  |  | Sex (M) | 0.24 | 0.2 | 1.16 | 0.247 |
|  |  | Recapture (Yes) | -0.05 | 0.26 | -0.21 | 0.833 |
|  |  | Sex (M):Recapture (Yes) | 0.21 | 0.3 | 0.69 | 0.494 |
| Activity (binomial) | **Males**<br>(161/101/21) | Intercept | 0.27 | 0.4 | 0.63 | 0.526 |
|  |  | Invasion zone (Edge) | -0.27 | 0.47 | -0.56 | 0.573 |
|  |  | Recapture (Yes) | 0.46 | 0.36 | 1.29 | 0.197 |
|  | **Females**<br>(129/98/23) | Intercept | -0.44 | 0.56 | -0.77 | 0.439 |
|  |  | Invasion zone (Edge) | 1 | 0.65 | 1.56 | 0.118 |
|  |  | Recapture (Yes) | 0.56 | 0.45 | 1.23 | 0.291 |
|  | **Edge**<br>(181/127/120) | Intercept | 0.068 | 0.35 | 1.94 | 0.052 |
|  |  | **Sex (M)** | **-0.78** | **0.37** | **-2.1** | **0.031** |
|  |  | Recapture (Yes) | 0.64 | 0.38 | 1.7 | 0.090 |
|  | **Source**<br>(109/75/7) | Intercept | -0.5 | 0.45 | -1.13 | 0.259 |
|  |  | **Sex (M)** | **0.91** | **0.42** | **2.14** | **0.032** |
|  |  | Recapture (Yes) | 0.25 | 0.44 | 0.575 | 0.565 |

